# A gradient of complementary learning systems emerges through meta-learning

**DOI:** 10.1101/2025.07.10.664201

**Authors:** Zhenglong Zhou, Anna C. Schapiro

## Abstract

Long-term learning and memory in the primate brain rely on a series of hierarchically organized subsystems extending from early sensory neocortical areas to the hippocampus. The components differ in their representational attributes and plasticity, with evidence for sparser, more decorrelated activity and faster plasticity in regions higher up in the hierarchy. How and why did the brain arrive at this organization? We explore the principles that allow such an organization to emerge by simulating a hierarchy of learning subsystems in artificial neural networks (ANNs) using a meta-learning approach. As ANNs optimized weights for a series of tasks, they concurrently meta-learned layer-wise plasticity and sparsity parameters. This approach enhanced the computational efficiency of ANNs, promoting hidden activation sparsity while benefitting task performance. Meta-learning also gave rise to a brain-like hierarchical organization, with higher layers displaying faster plasticity and a sparser, more pattern-separated neural code than lower layers. Early layers peaked early in their plasticity and stabilized, whereas higher layers continued to develop and maintained elevated plasticity over time, mirroring empirical developmental trajectories. Moreover, when trained on dual tasks imposing competing demands for item discrimination and categorization, ANNs with parallel pathways developed distinct representational and plasticity profiles, convergent with the distinct properties observed empirically across intra-hippocampal pathways. These results suggest that the macroscale organization and development of heterogeneous learning subsystems in the brain may emerge in part from optimizing biological variables that govern plasticity and sparsity.

## Introduction

To process and then learn from incoming information in the environment, the primate brain relies on a hierarchy of subsystems, which initiate in early sensory neocortical areas and converge on the hippocampus [1, 2]. Subsystems at different levels of this hierarchy exhibit very different properties, including in their plasticity — the degree of modifiability of neuronal responses by experience, and the nature of their representational coding — the range of inputs a neuron responds to [3, 4, 5, 6, 7]. Is there a broad structure and purpose that organizes the variation in these key properties across the brain?

Plasticity, which is ubiquitous in the brain, appears to increase along the neocortex-to-hippocampus hierarchy: early cortical areas often require at least several days of training to exhibit modified neuronal tuning, whereas higher-level visual areas can undergo more rapid changes [7, 8, 9, 10, 11]. Regions within the medial temporal lobe (MTL), and especially the hippocampus, in turn exhibit even faster plasticity, sometimes needing only one experience to drive lasting changes [3, 12, 4]. Moreover, there are differing developmental trajectories in this plasticity, with sensory neocortical plasticity peaking early in life and stabilizing thereafter, whereas higher-order regions reach peak plasticity later and retain it into adulthood [13, 14].

Representational sparsity appears to follow a parallel gradient across this hierarchy: activity in the medial temporal lobe (MTL) is considerably sparser than in neocortical regions [15], and within the MTL, sparsity increases from parahippocampal cortex to entorhinal cortex to hippocampus [16, 6, 17, 18], both in the proportion of neurons activated by a given input and in the number of distinct inputs that individual neurons respond to. In addition to these hierarchical gradients culminating in the hippocampus, plasticity and sparsity also vary across parallel pathways within the hippocampus itself [19, 20], suggesting that there is both serial and parallel macro-organization in these processing streams.

To understand how these systems support learning and memory, extensive theoretical work has sought to capture the properties and interactions across them [21, 22, 23, 24, 25, 26], or to explain why the brain requires multiple learning systems [27, 2, 28, 21]. One foundational account of neocortex–hippocampus interactions is the Complementary Learning Systems theory, which proposed an adaptive division of labor: the neocortex gradually builds overlapping, distributed representations that store our general understanding of the world, whereas the hippocampus rapidly creates sparse, pattern-separated codes that encode individual episodic memories [2, 29, 30].

While crucial in providing insight into the complementary functions of these different memory systems, this prior work modeling the hippocampus and neocortex has not provided an explanation for how these distinctions between multiple systems emerged in the first place. They have presupposed distinct modules with different parameters, effectively hard-coding their differences into the model architectures. And by modeling properties of the adult brain from the start, such models do not account for development phenomena like the earlier peak in plasticity in early sensory cortex relative to higher-order regions. Although some studies have investigated the benefits of heterogeneous plasticity in computational models [31, 32], they do not explain how plasticity varies across different brain areas. Finally, neither the hippocampus nor the neocortex is homogeneous, contrary to the original simple Complementary Learning Systems formulation; instead, both contain subregions with heterogeneous memory codes and varying degrees of plasticity. These limitations thus leave fundamental questions unresolved: why and how did multiple learning systems with heterogeneous properties emerge?

Here, we explore principles that allow a hierarchy of learning subsystems to emerge in artificial neural networks (ANNs), using a meta-learning approach. Many forms of ANNs, including convolutional neural networks and transformers, have demonstrated the capacity to develop through experience representations that mirror neural data in animals and humans [33, 34, 35, 36]. However, conventional approaches optimize weights but hard-code area hyperparameters, limiting the ability to explain the emergence of macro-organization across areas. To address this limitation, we enable ANNs to adaptively modulate layer-wise plasticity and sparsity, two key elements of across-area macro-organization, by extending a meta-learning method designed to minimize interference during continual learning [37]. Specifically, in our approach, as the network optimizes its weights for each task, it concurrently meta-learns layer-specific learning rates and layer-specific sparsity parameters that regulate within-layer competition, thereby allowing distinct plasticity and sparsity profiles to emerge across layers. Unlike many alternative methods for determining learning rates [38, 39, 40], the meta-learning approach affords the direct optimization of these variables for task performance [37, 41]. We assess both the impact of this approach on task performance and its tendency to produce a brain-like organization of heterogeneous learning systems.

We demonstrate that our meta-learning approach not only enhances the computational efficiency of ANNs but also promotes the emergence of brain-like distinctions reminiscent of the neocortex-hippocampus hierarchy. First, our models develop a sparse, efficient memory code that activates substantially fewer neurons while achieving equivalent or superior task performance compared to models that do not meta-learn layer-specific plasticity and sparsity parameters. Second, they form a brain-like hierarchical structure where higher layers develop greater plasticity, sparser representations, and enhanced pattern separation relative to lower layers. Third, the developmental trajectory of these networks mirrors that of the brain: the lowest layer rapidly achieves peak plasticity which then fades, whereas higher layers continue to adapt and remain plastic over time. Finally, our approach captures distinctions between the parallel intra-hippocampal pathways (i.e., the trisynaptic and monosynaptic pathways): in ANNs with pathways of different sizes trained on tasks with opposing demands of item recognition versus categorization, the larger pathway — akin to the trisynaptic pathway — updates weights more quickly, exhibits sparser activity, and plays a more important role in exemplar differentiation. Taken together, these results suggest that the organization of heterogeneous memory subsystems in the brain may arise through a learning process that tunes the biological variables that govern plasticity and activity levels in the service of optimized behavior.

## Results

### Meta-learning layer-wise plasticity and sparsity

We investigate how a hierarchy of learning subsystems can emerge in feedforward ANNs by concurrently optimizing both weights and two sets of per-layer variables *lr* and *s*, which govern layer-wise plasticity and sparsity, respectively. Through this approach, ANNs learn distinct plasticity profiles and memory codes across hidden layers. Specifically, *lr* represents layer-specific learning rates (Figure 1a), controlling how quickly weights at a particular level of the hierarchy are updated. *s* modulates layer-wise representational sparsity through a within-layer competition mechanism. The mechanism builds on the k-winners-take-all approach of Bricken et al. [42], which retains only the top k active neurons in a hidden layer. In each layer, the bias term is omitted, and the (k+1)-th highest activity is subtracted from every neuron’s output prior to applying the ReLU activation, ensuring that only the k most active neurons contribute to the final output. However, in this and related k-winners-take-all approaches [43, 42, 44, 45, 46], sparsity is fixed by the choice of *k*. To enable ANNs to learn and regulate sparsity explicitly, our approach introduces a learnable, layer-specific scalar parameter *s*. In each layer, the sparsity parameter *s*_*h*_ scales the *k*-th highest activity before subtraction, allowing the network to adjust the proportion of units with non-zero ReLU outputs. In practice, increasing *s*_*h*_ results in fewer active units, while decreasing it leads to more active units (Fig. 1b).

**Figure 1.**
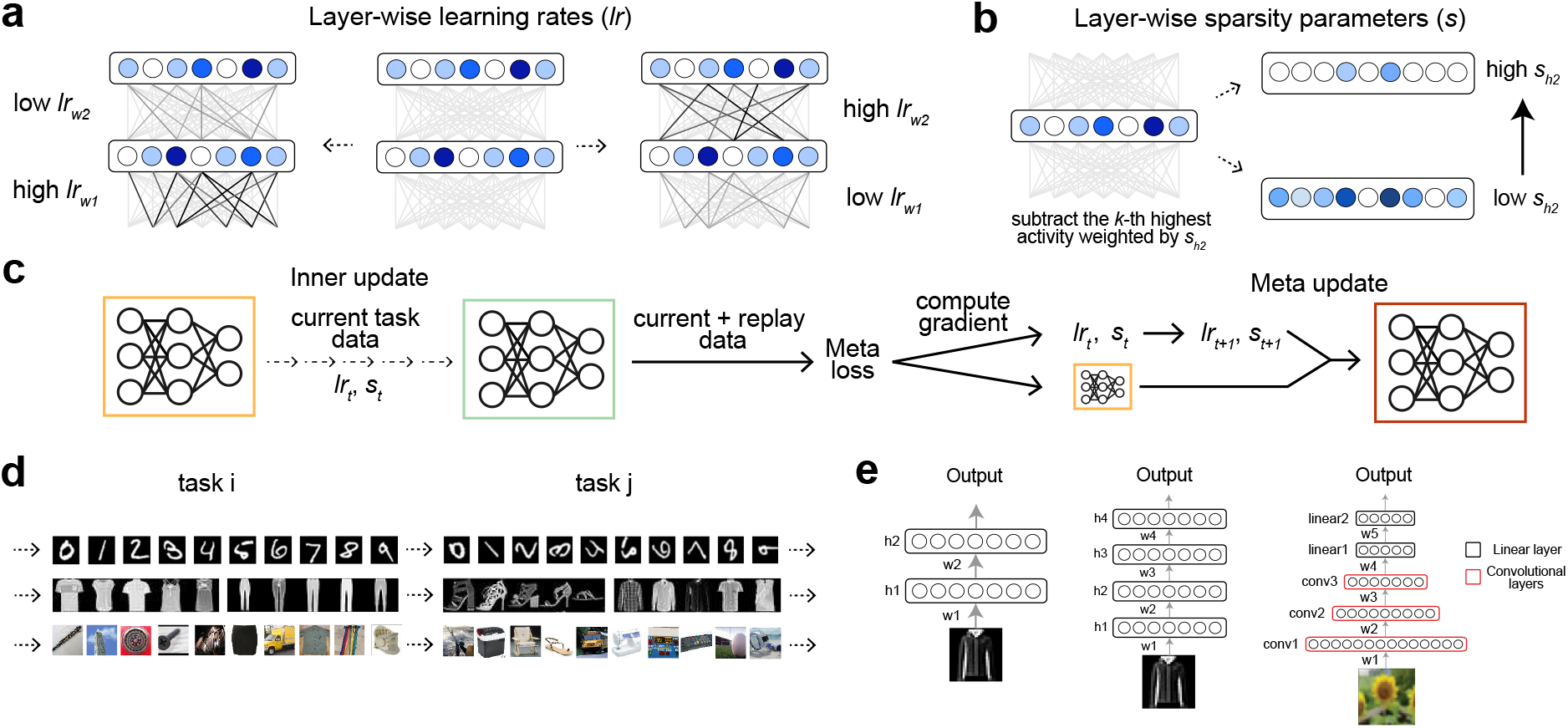
Meta-learning adaptive learning rates and sparse representations. **a**. The model meta-learns a distinct learning rate for each set of weights, allowing different layers within the hierarchy to adapt weights at varying rates. **b**. Each layer of the model meta-learns a per-layer parameter *s* that regulates representational sparsity via a within-layer competition mechanism. During the forward pass, the *k*-th highest unit activity is scaled by *s* and subtracted from all unit activities in that layer. By adapting *s*, the network dynamically controls the degree of sparsity in each layer. **c**. The meta-learning procedure alternates between inner updates and meta-updates. During the inner update, model weights are adjusted to optimize performance on data from the current task. Prior to the meta-update, a meta-loss is computed based on the performance of the inner-updated model on a mixture of current and past task data. Gradients from this meta-loss are then used to update the pre-inner-update weights, as well as layer-specific learning rates and sparsity parameters, enabling the model to adapt its learning dynamics and representations across layers. **d**. Experiments were conducted across multiple datasets, including rotated MNIST, Fashion MNIST, and CIFAR-100, to evaluate the generality of results across datasets. In each experiment, the model was exposed to a sequence of tasks introduced incrementally, with each task comprising a distinct subset of the dataset’s categories. **e**. Examples of architectures used in the experiments, which include standard fully connected networks — evaluated on both rotated MNIST and Fashion MNIST— and convolutional neural networks.

Our approach extends the meta-learning framework introduced by Gupta et al. [37] by incorporating mechanisms for regulating layer-wise plasticity and representational sparsity. Following Gupta et al. [37], we evaluate this approach in a continual learning setting, where the network sequentially learns tasks. This setting simulates real-world scenarios where environmental statistics shift over time and the inclusion of multiple learning systems have been theorized to be advantageous [2]. At each time-step *i* during training, the models optimize *θ*_*i*_ (model weights), *lr*_*i*_, and *s*_*i*_ for task *t* using a two-tiered process. First, the inner update applies a series of *j* weight updates on a batch of samples from task *t*, transforming 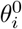 (identical to *θ*_*i*_) into 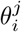. Next, the meta update evaluates 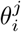 using a mixture of samples from task *t* and samples from a replay buffer of previous task data [37], computing a meta loss. Finally, the model backpropagates gradients of this meta loss with respect to 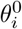, *lr*_*i*_, and *s*_*i*_, producing *θ*_*i*+1_, *lr*_*i*+1_, and *s*_*i*+1_. During the meta update, replay samples are drawn from a small memory buffer that accumulates data from previous tasks, facilitating gradient alignment between current and past tasks [47].

Our central question is whether the distinctions in plasticity and sparsity observed empirically along the sensory neocortex-hippocampus hierarchy emerge in ANNs that jointly meta-learn *lr, s*, and *θ*. As a baseline, we include models that meta-learn *θ* only, an approach that has matched or surpassed the performance of several established continual learning approaches [48, 49, 47]. We evaluate these models across multiple datasets, task settings, and model architectures (Fig. 1 d&e, Fig. 5 b), ranging from simple fully-connected ANNs assessed on basic benchmarks to convolutional neural networks evaluated on real-world classification tasks. In the following sections, we describe key results demonstrating that this approach not only improves computational efficiency but also induces layer-wise distinctions reminiscent of those observed along the neocortex-hippocampus axis.

### Enhanced computational efficiency with meta-learning

We first examined how meta-learning layer-wise learning rates (*lr*) and sparsity parameters (*s*) impacts task performance (i.e., test accuracy) and activation sparsity in ANNs, where sparsity is measured as the average proportion of non-zero hidden-unit activations across test samples. Models that meta-learn both *lr* and *s* achieve test accuracy equal to or exceeding that of baseline models across datasets (Fig. 2a), a result replicated over a broad range of commonly used learning rate initializations (Fig. A1&A3). This finding aligns with prior work showing that optimizing hyperparameters such as learning rate can improve ANN task performance [37, 50]. We also found that the models employ a sparse code, activating significantly fewer neurons than baseline models at test, which also replicated across different learning rate initializations (Fig. A2&A4). Unlike methods that directly optimize for sparsity (e.g., using regularization) [51], this approach achieves activation sparsity without optimizing an energy cost or sparsity objective. Together, these results show that meta-learning *lr* and *s* enhances the computational efficiency of ANNs, enabling them to achieve equal or superior test performance utilizing fewer active units.

**Figure 2.**
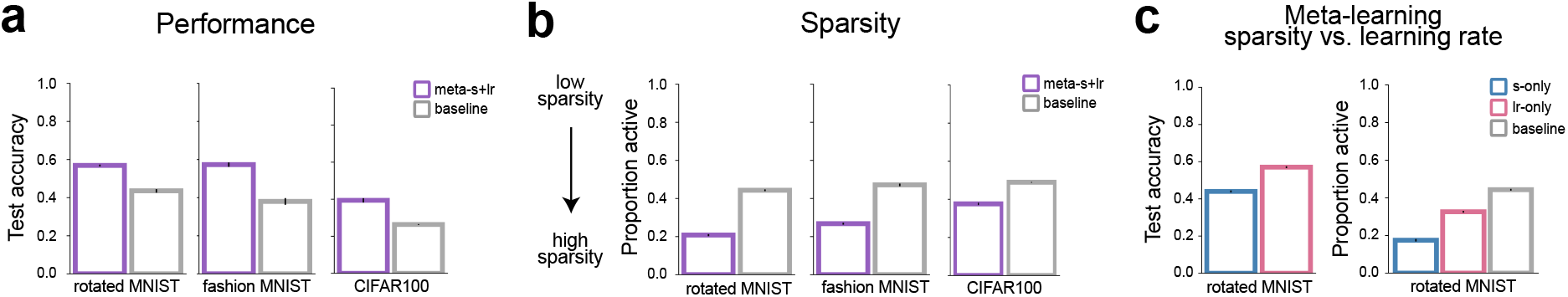
The impact of meta-learning learning rates and sparsity parameters on performance and representational sparsity. **a**. Models that meta-learn layer-specific learning rates and sparsity parameters consistently achieve higher test accuracy than baseline models across multiple datasets, demonstrating the benefits of adaptive optimization of *lr* and *s*. Error bars reflect ±1 standard error of the mean across independent model runs initialized with distinct random seeds. **b**. Models with meta-learned learning rates and sparsity parameters exhibit greater activation sparsity compared to baseline counterparts across multiple datasets, suggesting the emergence of more efficient task-specific representations. **c**. Meta-learning of learning rates primarily drives performance gains, while meta-learning of sparsity parameters contributes more strongly to increased activation sparsity.

To isolate the contributions of meta-learning *lr* and *s* to sparsity and performance, we examined models that meta-learned only one of the two (i.e., *lr*-only or *s*-only). Both *lr*-only and *s*-only models exhibit greater sparsity than baseline models at test, suggesting that both encourage the formation of sparse representations. *s*-only models achieve even greater sparsity than *lr*-only models, while *lr*-only models attain higher task accuracy (Fig. 2c). The two variables thus have complementary benefits for computational efficiency, with meta-learning *lr* doing more to enhance task performance (achieving high performance with less training) and meta-learning *s* doing more to promote activation sparsity.

### The emergence of a brain-like hierarchy through meta-learning

Our results thus far show that meta-learning *lr* and *s* facilitates the computational efficiency of ANNs. We now turn to the question of whether meta-learning *lr* and *s* leads to differentiation of plasticity and neural coding across hidden layers in ANNs. We evaluate whether meta-learning induces an organization in ANNs that aligns with that observed in the neocortex-hippocampus hierarchy, including faster plasticity and sparser neural activity in higher-level subsystems.

We first examine these properties in shallow ANNs with two hidden layers (Fig. 1e left). Through meta-learning, a hierarchical organization of layer-wise learning rates emerges in these networks: *lr*_*w*2_ (i.e., the learning rate for *w*2, Fig. 1e left) becomes substantially higher than *lr*_*w*1_ (Fig. 3c) across model runs with different random seeds and layer sizes (Fig. A5b). This pattern gradually emerges over the course of training: Early on, *lr*_*w*1_ increases rapidly and surpasses *lr*_*w*2_; as training progresses, *lr*_*w*1_ plateaus, decreases, and eventually falls below the level of *lr*_*w*2_ (Fig. 3f). The developmental trajectories suggest that the model learns to quickly fine-tune *h*1 representations initially, then gradually stabilizes them while increasingly adapting *h*2 representations for later learning. The non-monotonic development of *lr*_*w*1_ and *lr*_*w*2_ suggests that meta-learning does not simply organize learning rates by the proximity of their weights to the output.

**Figure 3.**
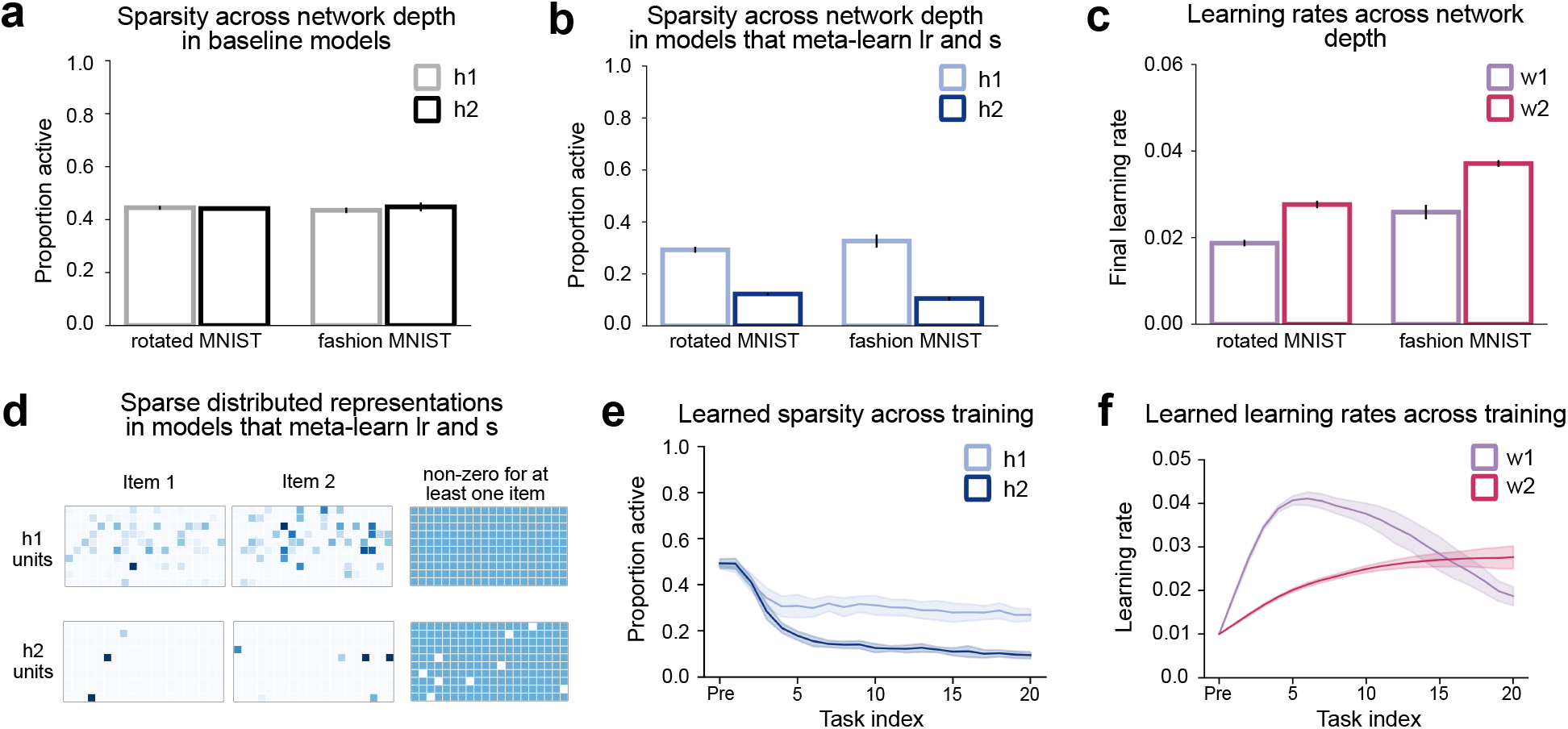
Emergence of a hierarchical organization of learning rates and sparsity in shallow ANNs. **a**. In baseline models that meta-learn only weights, hidden layers exhibit uniform levels of activation sparsity, lacking differentiation across layers. **b**. In two-layer models that meta-learn both learning rates and sparsity parameters, the upper hidden layer exhibits greater activation sparsity, suggesting the emergence of a hierarchical organization of sparse representations. **c**. Models that meta-learn both learning rates and sparsity parameters consistently assign higher learning rates to the top hidden layer, reflecting a hierarchical tuning of plasticity across layers. **d**. Examples illustrate greater activation sparsity in the top hidden layer (h2) compared to the first hidden layer (h1). Although most units in h2 are inactive for any given input, the majority are active for at least one item across the dataset, consistent with the formation of sparse distributed representations. **e**. Early in training, activation sparsity decreases at a similar rate in both hidden layers (h1 and h2). Over time, however, sparsity levels diverge, with h2 showing a more pronounced decline, indicating layer-specific adaptation dynamics in representational sparsity. **f**. The learning rate for weights in the first hidden layer (w1) increases more rapidly than that of the second hidden layer (w2) early in training, then stabilizes, and gradually declines, eventually falling below the learning rate assigned to w2. This pattern indicates a dynamic reallocation of plasticity across layers over the course of learning.

A hierarchical organization of activation sparsity concurrently emerges in this simple network. We measure activation sparsity for each hidden layer as the average proportion of active units across test samples. Early during training, *h*1 and *h*2 show increasing levels of sparsity with little differentiation. As learning progresses, *h*2 sparsity stabilizes, while *h*1 sparsity continues to increase (Fig. 3e). This leads to a hierarchy of activation sparsity, with the higher layer *h*2 showing substantially higher sparsity, a result that held across simulations with different layer sizes (Fig. A5a) and across tasks (Fig. 3b). In contrast, baseline models without meta-learning of *lr* and *s* show no hierarchical differentiation of sparsity between hidden layers (Fig. 3a).

We also find that *h*2 activity shows greater pattern separation across tasks than *h*1 activity (Fig. A6a). To quantify pattern separation across tasks for each layer, we extracted the activation vector evoked by each task at test and computed the mean pairwise correlation between activation vectors from distinct tasks, where lower mean correlations indicate stronger pattern separation (see Methods). This reveals lower across-task correlation in *h*2 activation compared to *h*1 at test, suggesting that *h*2 develops a greater capacity to de-correlate inputs along with increased sparsity through experience. Note that high sparsity does not necessarily imply high pattern separation, as a network activating the same small subset of units for all tasks can be sparse without achieving pattern separation. The co-development of sparsity and pattern separation suggests that these networks form sparse distributed representations [52, 53], where a small subset of units activate for individual inputs, yet nearly all units contribute to the representation of at least one input. These properties of sparse distributed representations are evident in both hidden layers (Fig. 3d), consistent with a previous finding that meta-learning facilitates their formation [54].

Next, we assess whether these patterns generalize to deeper and more complex network architectures. In deeper networks with four hidden layers (Fig. 1e middle), meta-learning *lr* and *s* leads to a similar hierarchical pattern, where the top hidden layer exhibits higher learning rates (Fig. 4c), increased sparsity (Fig. 4a), and enhanced pattern separation across tasks (Fig. A6b). These networks exhibit similar developmental trajectories, with the learning rate for the lowest hidden layer’s incoming weights increasing most rapidly early on (Fig. 4d), while sparsity stabilizes faster in the lower hidden layers (Fig. 4b). Similar hierarchical organizations of learning rates and sparsity emerge in convolutional neural networks (Fig. 1e right) that meta-learn *lr* and *s* for real-world image classification (Fig. 1d bottom row). Notably, however, the learning rates in the lower layers do not exhibit a strictly monotonic increase (Fig. 1c&f). In sum, we observe that a hierarchical organization of learning rates and sparsity consistently emerges across a wide range of model architectures, datasets, and hyperparameters through meta-learning. The hierarchical organization that emerges in our networks mirrors key features of the neocortex-hippocampus axis. Specifically, higher-level structures exhibit faster plasticity, higher activation sparsity [16, 6], and more pronounced decorrelation of task inputs [17] compared to lower regions. The differential development of these properties across layers also mirrors the brain, where early cortex develops peak plasticity early in life and stabilizes, while higher-order regions peak later and retain greater plasticity into adulthood [13]. Thus, meta-learning variables that modulate plasticity and sparsity offers an account of the development of distinct subsystems across the neocortex-hippocampus hierarchy.

**Figure 4.**
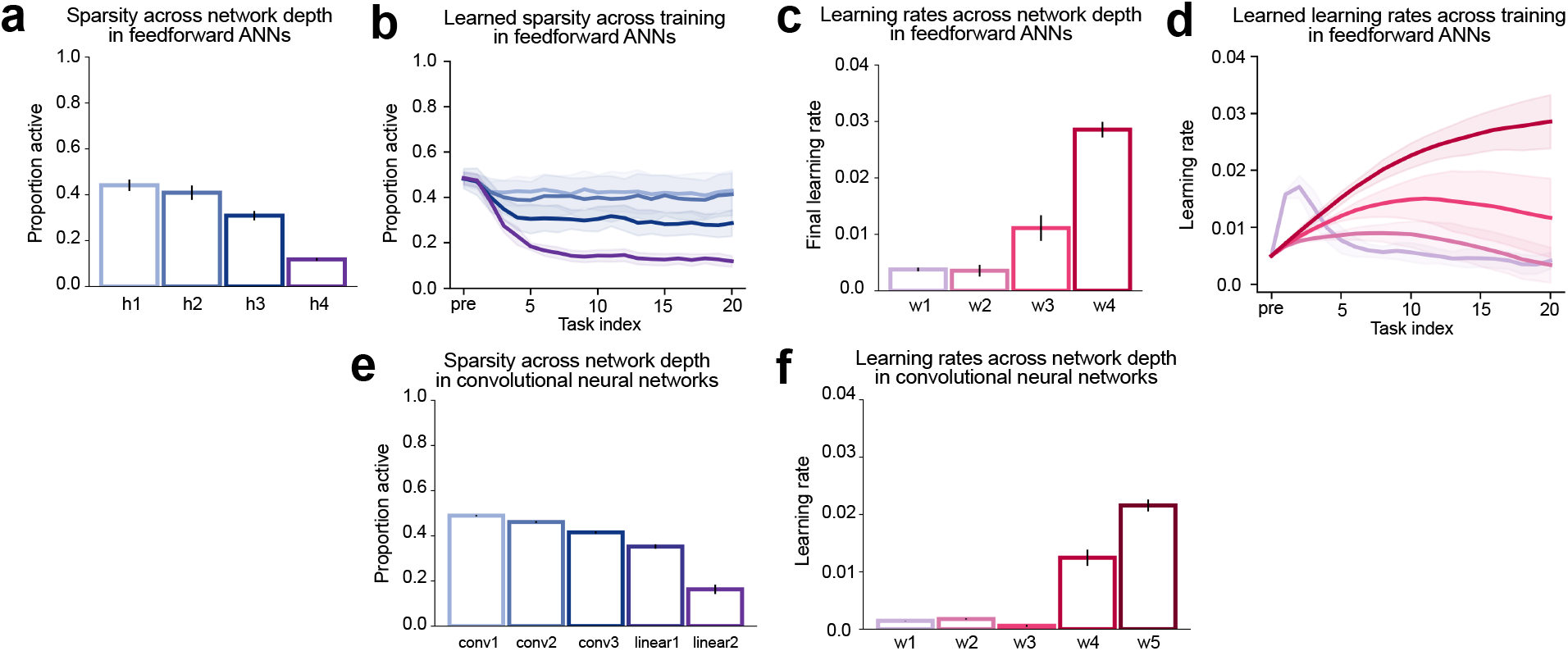
Generalization of hierarchical patterns of learning rates and sparsity in deeper ANNs. **a**. In standard feedforward ANNs with four hidden layers, meta-learning induces a hierarchical organization of activation sparsity, with the highest sparsity observed in the topmost layer. **b**. This differentiation in sparsity across layers emerges early in training. **c**. A similar hierarchical structure develops in the learning rates of meta-learned models, with the final layer (w4) acquiring the highest learning rate. **d**. The developmental trajectories of learning rates follow a consistent pattern: after a sharp initial rise, w1 declines, while w4 continues to increase, establishing a hierarchical structure. **e**. A comparable hierarchy of activation sparsity also emerges in convolutional neural networks. **f**. Likewise, convolutional neural networks exhibit a structured hierarchy of learning rates under meta-learning.

### Explaining the origin of distinct intra-hippocampal pathways

Thus far, we have shown that meta-learning enables ANNs to capture hierarchical distinctions across sub-systems in the neocortex-hippocampus hierarchy. In the brain, these properties also vary across parallel pathways. Within the hippocampus, two parallel anatomical pathways — the monosynaptic (MSP) and trisynaptic (TSP) pathways — process inputs from the entorhinal cortex (EC). The larger TSP exhibits sparser activity [5], greater pattern separation of task inputs [19, 55], and faster modulation of neural tuning with new experience [56, 57]. Furthermore, evidence suggests that the TSP is uniquely important for tasks that require discriminating individual exemplars [58, 59, 60]. To capture these distinctions, models of the hippocampus have constructed parallel pathways with different learning rates and sparsity [20, 61]. Here, we explore whether ANNs with parallel pathways develop these differences through meta-learning.

We explore ANNs with two parallel pathways at the top of a series of hidden layers (Fig. 5b), representing the two intra-hippocampal pathways sitting on top of the neocortex-hippocampus hierarchy. Following prior approaches, the two parallel pathways in the networks differ in size, with the larger pathway representing the TSP and a size ratio that loosely mimics that in the hippocampus [5]. We adopt a simplified architecture that allows independent contributions of the two pathways to the output, as opposed to both pathways running through a common CA1. There is evidence for differential topology of EC vs. CA3 inputs to CA1, which would provide a degree of this kind of independence [62, 63, 64]. To test the contribution of the two pathways to distinct task demands [20, 65], we train the model on a dual task [66] requiring concurrent mapping of each MNIST image to a category label (0-9) and an item-specific label (Fig. 5a). The task imposes opposing demands of categorization and exemplar discrimination, as it requires mapping items of the same category to a shared category label while assigning them unique item labels. The model optimizes *lr, s*, and *θ* as it meta-learns a series of dual tasks. This set of simulations allows us to test whether the meta-learning approach can account for the observed differences between parallel hippocampal pathways.

**Figure 5.**
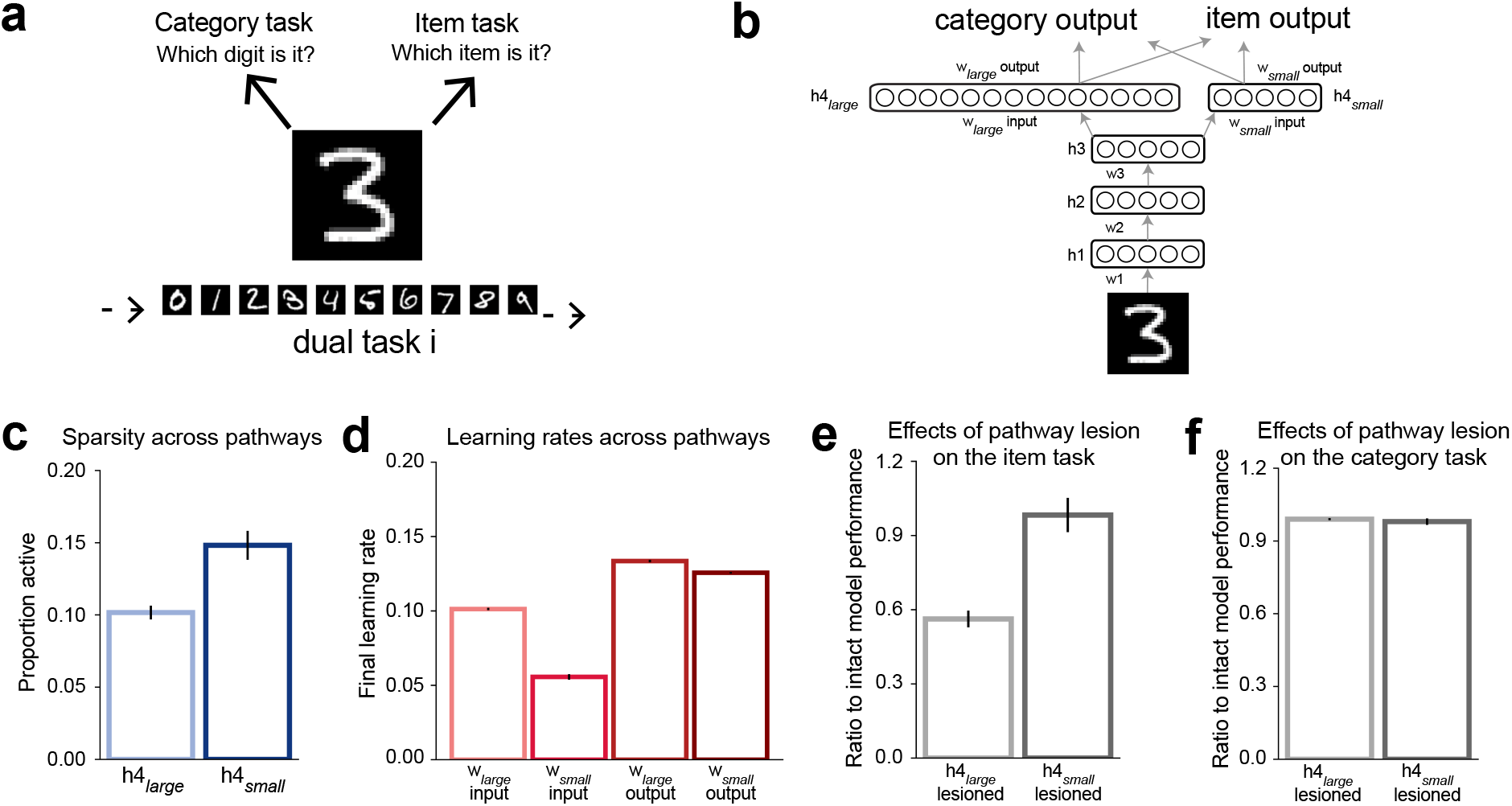
Differentiation of parallel pathways in ANNs through meta-learning of dual tasks. **a**. Models were trained on a series of dual tasks using rotated MNIST images, each requiring the classification of both category labels (0-9) and item-specific labels. **b**. The ANN architecture features two parallel pathways, mirroring intra-hippocampal pathways, with independent projections to the output. **c**. Following meta-learning of layer-specific learning rates and sparsity parameters, the larger pathway (*h*4_*large*_) develops higher activation sparsity than the smaller pathway. **d**.The same model assigns higher learning rates to weights in the larger pathway (*w*_*large*_ for input and output) compared to those in the smaller pathway. **e**. At test, disabling the larger pathway selectively impairs item-specific label identification more than disabling the smaller pathway. **f**. Disabling either pathway at test results in comparable performance on the category label task.

Through meta-learning, the two-pathway ANN develops hierarchical and parallel differentiation of learning rates and sparsity across its hidden layers. Replicating observations from single-pathway models in previous simulations, the two-pathway ANNs develop higher learning rates and increased activation sparsity in the top hidden layer. In addition, meta-learning gives rise to the emergence of distinct properties between the two pathways: The larger pathway develops higher learning rates (Fig. 5d), sparser task representations (Fig. 5c), and enhanced pattern separation of task inputs (Fig. A6c). These characteristics are reminiscent of differences observed between intra-hippocampal pathways [5, 19, 57]: The higher learning rates in the larger pathway in our model facilitate faster plasticity, akin to the role of the TSP in rapid encoding, and increased sparsity and greater pattern separation in the larger pathway align with the TSP’s known specialization in reducing overlap between inputs. Furthermore, we observed distinct contributions of the two pathways to task demands. At test, the larger pathway is more critical for exemplar discrimination: Disabling it impairs the model’s ability to identify item-unique labels more than disabling the smaller pathway (Fig. 5e). This functional differentiation aligns with empirical evidence that tasks requiring exemplar discrimination rely more heavily on the TSP [58, 59, 60]. In contrast, disabling either pathway results in comparable performance on the categorization task (Fig. 5f), suggesting that the two pathways develop redundant representations that support categorization. Together, these findings suggest that the two-pathway meta-learning model captures distinct computational properties consistent with the proposed division of labor within the hippocampus [20]. The meta-learning approach thus offers a computational framework for understanding the development of parallel — in addition to serial — circuits that contribute to complementary learning and memory processes.

## Discussion

How did a hierarchy of learning subsystems with distinct computational properties in the primate brain come to be? Instead of the traditional modeling approach of setting these properties manually to build architectures that resemble the brain, here we allow these properties to be learned and then evaluate whether they endogenously emerge in a manner that resembles the brain. If so, this would suggest that these properties are likely to be the result of learning processes that optimize these subsystems for adaptive behavior. First, we show that ANNs that meta-learn layer-wise plasticity and plasticity parameters develop a computationally efficient, sparse neural code, achieving equal or better performance with fewer active units than baseline models. Second, a brain-like hierarchical organization emerges in these models: Higher layers learn faster weight updates, exhibit sparser activity, and achieve greater input de-correlation than lower layers. Third, the models capture patterns of brain plasticity development, with an early peak and gradual reduction in lower layers while higher layers develop and maintain plasticity through subsequent experience. Finally, we show that this approach also accounts for differences in plasticity and sparsity between parallel pathways within the hippocampus: ANNs with parallel pathways of different sizes, trained on tasks requiring item recognition and categorization, develop divergent properties similar to intra-hippocampal pathways, with the larger pathway exhibiting higher learning rates, sparser activity, and a critical role in distinguishing exemplars. Together, our approach captures the heterogeneous properties observed across hippocampal and neocortical regions, and suggests that the optimization of biological variables modulating plasticity and sparsity may underlie the brain’s organization and development of multiple learning subsystems.

Much empirical work has investigated variations in plasticity, sparsity, and pattern separation across the neocortical-hippocampal hierarchy in adult primate brains, with a particular emphasis on the visual system. First, plasticity varies along this hierarchy. In perceptual learning tasks, changes in the selectivity and responsiveness of neurons in higher-level cortical areas like V4 and IT [7, 8] tend to require less training than those in V1 and V2 [67, 11, 9]. Compared to sensory neocortical areas, the MTL [68, 10] and the hippocampus in particular support much faster plasticity, such that individual MTL cortical neurons can develop novel response properties within just a few trials [3] and the hippocampus can develop new place fields after a single experience [4]. Second, activation sparsity appears to be hierarchically structured. There is evidence that, within the neocortex, sparsity increases progressively across the ventral visual stream [69] (although see [70]). MTL cortex and hippocampus exhibit much sparser activity than cortical sensory areas [6, 71, 72]. Sparsity also differentiates between MTL cortex and the hippocampus: the entorhinal cortex (a MTL input region to the hippocampus) displays sparser activity than the hippocampus in spatial tasks [16]. Furthermore, these properties differ between the two intra-hippocampal pathways, with the larger trisynaptic pathway exhibiting sparser activity [5], greater pattern separation of different inputs [19], and faster changes in neural responses to novel information [57]. Overall, these observations support theories suggesting that plasticity, sparsity, and pattern separation are organized hierarchically in the brain [73, 74, 2, 7, 75]. Our results suggest that this organization is optimized for behavior.

Canonical memory theories have put forward a two-system view [2, 76] in which the hippocampus and neocortex execute distinct and complementary computations. However, McClelland et al. [2] offered the speculation that plasticity may in fact be hierarchically organized, with rapid learning in the hippocampus, slow learning in sensory neocortex, and intermediate rates in regions surrounding the hippocampus. We find evidence for this more graded view, finding hierarchies of complementary learning systems. Other work has provided consonant views about the hippocampus and neocortex lying along a graded hierarchy of this kind [77, 74, 75]. Clarifying the precise nature of this gradient will require further empirical investigation. For example, although the disparity in sparsity between the neocortex and medial temporal lobe is well documented, studies examining the variation of sparsity across neocortical subregions have produced mixed results [69, 70], underscoring the need for additional work. Moreover, it remains to be determined whether analogous organizational principles might extend to other sensory domains, such as auditory processing.

We found in our simulations, as in the brain, that higher sparsity tends to co-occur with faster learning rates. We view these not as parallel, independent gradients that happen to align but rather as functionally interrelated properties. Distributed representations, where the same computational units are involved in representing many related memories [78], support efficient memory representation and powerful generalization, but require slow, interleaved learning in order to avoid new memories overwriting old ones that make use of the same weights and units [2]. Sparse, pattern-separated representations, in contrast, are less efficient and less adept at generalization but have the complementary strength of avoiding interference between similar memories; this representational strategy uses largely independent sets of units for each memory, which affords rapid learning by avoiding the potential for overwriting. Sparse representations thus allow for the possibility of rapid learning while distributed representations require slow learning, leading to the natural covariation of these properties.

What biological processes and variables might implement such a meta-learning mechanism in the brain? While our meta-learning process unfolds on the same timescale as task learning, analogous processes may operate over much longer developmental or evolutionary timescales in the brain. Developmentally, two key factors have been proposed to regulate the formation of sparsity and plasticity in neural circuits. First, the maturation of inhibitory circuits has been proposed to drive the sparsification and decorrelation of neural activity [79]. Second, progressive myelination gradually restricts synaptic plasticity and closes critical periods [80, 81]. These two factors follow distinct maturational schedules: neural inhibition reaches functional maturity relatively early, between late childhood and adolescence, and exhibits only modest timing differences between sensory and higher association areas [82, 83, 84]. In contrast, myelination continues into the late twenties or even thirties and shows much larger variation in its developmental schedule across areas [85, 86, 87, 88]. These developmental timelines mirror dynamics in our meta-learning models: network sparsity across layers stabilizes early, with only slight variation in when each layer stabilizes, whereas learning rates continue to adjust over longer periods across layers. These factors have been proposed to serve multiple roles. For instance, the maturation of parvalbumin-positive (PV) inhibitory neurons has been proposed to regulate the timing of critical-period plasticity peaks across cortical regions, consistent with its hierarchical development in the monkey visual cortex [13, 14]. Moreover, both inhibitory circuit maturation and myelination are experience-dependent [89, 13], making it possible that the brain hierarchically calibrates these factors to optimize sparsity and plasticity for behavior during development [90]. Another (not mutually inconsistent) possibility is that an analogous form of meta-learning occurs across evolutionary timescales, where an optimized hierarchy or developmental schedule of plasticity across neural systems may have evolved in parallel with the optimization of brain architecture.

Modern deep learning systems are orders of magnitude more energy-intensive than the human brain [51, 91], which achieves exceptional computational efficiency through mechanisms such as sparse coding [92]. Addressing this disparity is a key focus in machine learning research, with efforts dedicated to reducing energy consumption while preserving performance of the systems. A common approach is to increase sparsity of activation in these models, through techniques such as weight pruning and regularization. Building on these approaches, our work demonstrates the potential of a meta-learning framework to promote optimally sparse activation. Rather than explicitly optimizing for sparsity, our method focuses on maximizing task performance, allowing sparsity to emerge naturally. Our simulations demonstrate that this approach not only enhances activation sparsity but also maintains strong task performance. Our work suggests that the metalearning approach provides a complementary strategy for reducing the energy demands of artificial systems and may offer insights into the design of brain-like neural architectures.

Future work could generalize this meta-learning approach to capture other kinds of systematic variations across memory systems and test whether these differences reflect optimized memory processing. For instance, different subfields of the hippocampus have different degrees of recurrence, with high recurrence in CA3 in particular thought to support pattern completion [76]. Existing models hard-code this architectural feature, but a meta-learning approach could explore whether it emerges in the process of optimization. Another use could be in exploring the role of different sleep stages in memory consolidation [93], where stochasticity and strength of communication between memory systems across sleep stages could be optimized instead of hard-coded [24].

In conclusion, our work provides evidence that meta-learning layer-specific plasticity and sparsity enables ANNs to develop properties that mirror the graded organization of learning systems along the neocortex–hippocampus axis. Our models not only develop an efficient, sparse memory code but also recapitulate structured variations in plasticity and sparsity across both serial and parallel processing streams, along with developmental trajectories mirroring those observed in the brain. By allowing these properties to emerge rather than hard-coding them, our approach goes beyond conventional two-system models and accounts for the heterogeneous computational features of hippocampal and neocortical regions. These findings support the notion that optimization processes—whether during development or across evolutionary timescales—may have finely tuned these learning systems for adaptive behavior. This meta-learning approach could serve as a flexible framework for exploring systematic variations across neural systems and patterns of brain development across humans and other species, and may inform the design of computationally efficient, brain-inspired machine learning architectures.

## Methods

### Model architectures

All models share a common feedforward architecture comprising an input layer, one or more hidden layers, and an output layer. To address distinct computational tasks, these networks are instantiated in different forms, including fully connected, convolutional, and dual-pathway models. Hidden layers omit bias terms and use the ReLU activation function. Each layer maintains its own learnable learning-rate *lr* and sparsity *s* parameters, enabling fully flexible meta-learning of both plasticity and activation properties.

Each layer has independently tuned learning rates (*lr*) and sparsity parameters (*s*). This approach enables flexible meta-learning, where each set of weights and each layer can adjust its dynamics autonomously. Each set of weights is equipped with a layer-specific learning rate *lr*, which governs the speed of weight updates. Each layer has a layer-specific sparsity parameter, *s*, that dynamically regulates the proportion of active neurons. The sparsity mechanism builds on a k-winners-take-all approach [42]: During the forward pass, unit activities are adjusted by subtracting the (*k* + 1)-th highest activity value before applying the ReLU function, thereby retaining only the top *k* activations. Our mechanism extends this approach by introducing a layer-specific parameter *s* which multiplies the *k*-th highest activity *k*_*a*_ of the layer before subtracting it from all activations. For each layer, *s* was initialized to 0.01, and *k* was set to one percent of the layer size, rounded to the nearest integer. As a result, the network can flexibly modulate activation sparsity, such that higher values of *s* yield fewer units being active, and lower values allow for more active units. This flexible regulation of activation sparsity differs from previous models that enforce a fixed level of sparsity.

For fully-connected networks, we experimented with two variants to probe how meta-learning interacts with depth, testing hidden-layer sizes of 100, 200, 400, and 800 units. In the shallow variant, the network has two hidden layers, while the deeper variant has four hidden layers. In both cases, each hidden layer omits bias terms to simplify the dynamics of within-layer competition in accordance with Bricken et al. [42].

For convolutional neural networks (CNNs) that perform real-world image classification tasks, the architecture begins with three convolutional layers, each employing 160 filters of size 3 × 3 with a stride of 2 and a padding of 1. Each convolutional layer is immediately followed by a ReLU activation. After the input is processed by the convolutional layers, the feature maps are flattened and then processed by a series of fully connected layers: An initial linear layer reduces the dimensionality to 640, followed by a ReLU activation; a second linear transformation maintains a 640-dimensional representation with another ReLU; and a final linear layer projects the features onto the output. Throughout the CNN, layer-specific *lr* and *s* parameters are applied to both convolutional and fully connected layers.

For dual-pathway models that perform the dual tasks, the network first processes inputs through three hidden layers, each containing 200 units. The network then splits into two parallel pathways: a larger pathway (denoted by 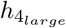) with 1000 units, and a smaller pathway 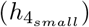with 200 units. The relative size ratio mirrors that between the trisynaptic pathway and monosynaptic pathway in the hippocampus [5]. Both pathways receive inputs from the third hidden layers and are fully connected to two distinct output layers corresponding to dual task demands, one for the category output and one for the item-specific label output for each image. Each layer in both pathways adapts its own *s* parameter to regulate activation sparsity. Moreover, weights of connections linking each path to the third hidden layer and to each of the output layers are governed by separate learning rates, allowing the meta-learning approach to independent optimize the plasticity of the two pathways.

### The meta-learning approach

Our meta-learning approach extends Gupta et al. [37]. In Gupta et al. [37], each task is first adapted in an inner loop and then evaluated on a meta loss. The meta-learning updates jointly optimize the model’s weight parameters *θ* and per-parameter learning rates. Here, we extend the method by meta-learning weights *θ* as well as layer-specific 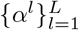and 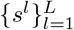. For brevity, we write 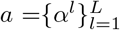and 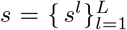to denote the set of all layer-specific parameters, and *α*^*l*^ and *s*^*l*^ to refer to parameters at layer *l*.

#### Inner update of model parameters

Given a new task *T* with training loss ℒ_*T*_ (*θ, s*), the inner-loop updates are performed separately for each layer *l*’s incoming weights *θ*^*l*^ over *K* steps. We denote the incoming weights of layer *l* after the *k*-th inner update as 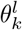, with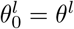. For *k* = 1, …, *K*, the update is given by:

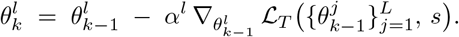

We denote the final adapted weights as *θ*_*K*_. During this inner-loop update, both *α* and *s* are held fixed.

#### Meta update of *a, s*, and *θ*

After the inner-loop updates, a meta loss is computed on a batch comprising samples from prior tasks as well as task *T* :

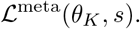

The overall meta-objective is to minimize this loss with respect to the model parameters *θ*, layer-specific learning rates *α*, and layer-specific sparsity parameters *s*

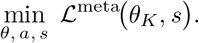

The formulas below are derived under the assumption *K* = 1; for *K >* 1, the corresponding gradients are obtained by backpropagating the meta-loss through each of the *K* inner update steps.

The meta-gradient for *α*^*l*^ is:

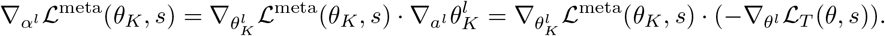

where 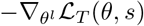 is the gradient of of the inner-loop update with respect to 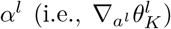. Using a meta learning rate *γ* for the per-layer learning rates, the update for each *α*^*l*^ is:

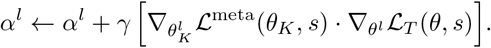

The meta-gradient for *s*^*l*^ is given by:

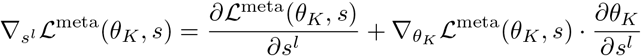

Since

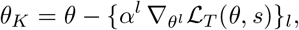

the derivative with respect to *s*^*l*^ is:

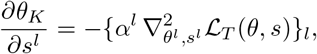

where 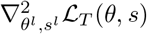 denotes the mixed partial derivatives with respect to *θ*^*l*^ and *s*^*l*^. Thus, the meta-gradient for *s*^*l*^ becomes:

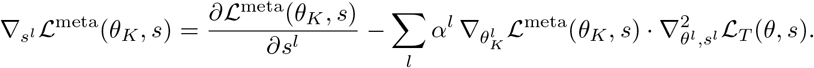

With a meta learning rate *δ* for *s*, its update rule is:

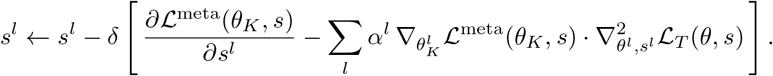

For each layer *l*, the meta-gradient with respect to *θ*^*l*^ is obtained by differentiating the meta loss through the inner-loop update:

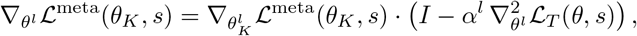

where *I* is the identity matrix and 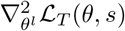 is the Hessian with respect to *θ*^*l*^. With the updated *α*^*l*^, the update is:

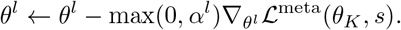

We set *γ* to 2*e*– 4 for single-pathway fully-connected networks trained on rotated MNIST, 5*e*–4 for single-pathway fully-connected networks trained on Fashion MNIST, 2*e*–5 for convolutional architectures, and 2*e*–2 for dual-pathway models; and we set *δ* to 2e-2, 5e-4, and 2e-1 for those same architectures, respectively. We assessed task performance and representational sparsity across a wide spectrum of initial learning rates *α*_0_ (Figs. A1–A4). Crucially, varying *γ* and *δ* over several orders of magnitude did not affect our main pattern of interest: higher hidden layers consistently learned more rapidly and developed sparser representations. However, to reveal these layer-wise, non-monotonic learning-rate dynamics, *γ* must be sufficiently large; an extremely small *γ* would prematurely truncate the learning curves (Fig. 3f) and mask these patterns.

All simulations reported here were built on the publicly available implementation from Gupta et al. [37], available at https://github.com/schapirolab/meta-CLS, using their default hyperparameter settings wherever possible. We note, however, a minor discrepancy between the methods as described in their paper and code implementation. Specifically, the manuscript [37] specifies that the meta-objective is evaluated only once, after the final inner-loop update, while their code computes the meta-loss at every inner-loop step. To ensure this did not impact our conclusions, we implemented both variants and found that they produced qualitatively identical results in terms of the observed layer-wise differences and comparisons to baseline models.

### Datasets and Task Paradigms

#### Benchmark Datasets

To evaluate the proposed meta-learning framework, experiments were conducted on three benchmark datasets: rotated MNIST, Fashion MNIST, and CIFAR-100. These datasets provide a range of visual recognition challenges, allowing assessment of the approach’s generalizability. Fashion MNIST and rotated MNIST were used for experiments with fully connected networks, while CIFAR-100 was employed in conjunction with convolutional neural networks (CNNs).

### Experiment Setup

The experiments employed a sequential, continual task-learning setting, simulating learning in non-stationary environments. In the rotated MNIST experiment, a sequence of 20 tasks was constructed, each corresponding to a different degree of rotation applied to the standard MNIST dataset images. For the Fashion MNIST dataset, five tasks were created, each consisting of two categories from the full dataset. In the CIFAR-100 experiment, twenty tasks were defined, where each task included five distinct categories from the dataset.

Across all experiments, tasks were introduced sequentially, ensuring that the model encountered new data incrementally and without direct access to previously learned tasks. To mitigate catastrophic forgetting, a replay buffer was incorporated which maintained a subset of previously encountered samples, following the approach of Gupta et al. [37]. This setup imposed constraints on the learning process, simulating real-world scenarios where knowledge is acquired progressively in a sample-efficient manner. Each task in the rotated MNIST, Fashion MNIST, and CIFAR-100 contained 200, 200, and 250 samples, respectively.

#### Design of the Dual-Task

To examine whether the proposed meta-learning framework can account for differences between intra-hippocampal pathways, we adopted a dual-task paradigm inspired by Kang and Toyoizumi [66], assessing the network’s ability both to recognize individual items and to integrate shared features across related items. These functions align with key computational roles attributed to the hippocampus, as suggested by previous models positing that distinct subsystems within the hippocampus support complementary processes: one extracting common structure across experiences, while another performs pattern separation to preserve individuated item representations [65].

The dual-task paradigm was implemented as an extension of the rotated MNIST experiment. In addition to the standard classification task, each image was assigned a unique item-specific label, requiring the network to learn both item categorization and discrimination. This augmentation introduced 2000 distinct item-specific labels across the dataset, significantly increasing the representational demands of the task. By integrating both objectives within a single model architecture, the dual-task provided a stringent test of the network’s ability to learn under competing task demands.

### Pattern separation analysis

We assessed pattern separation at each hidden layer using representational similarity analysis (RSA; Fig. A6). For each random initialization, we extracted post-training activation patterns for the 20 rotated MNIST tasks at each hidden layer *𝓁* ∈*{*1, …, *L}*, yielding matrices of dimension (trials ×units). At each layer and for every ordered task index pair (*i, j*), we randomly drew *m* activation vectors (without replacement) from each task and computed Pearson correlations for all *m*^2^ cross-condition vector pairs. The mean correlation defined the RSA element 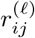. We then collapsed the full 20 × 20 RSA matrix to a scalar pattern-separation metric *s*^(*𝓁*)^ by averaging over all entries—lower values indicating stronger representational orthogonalization. Comparing *s*^(*𝓁*)^ across layers thus reveals how pattern separation among task representations vary across the network hierarchy.

## Acknowledgments

We thank Monami Nishio for valuable discussions about this work.

## Code Availability

The code used to run simulations in this study is available at: https://github.com/schapirolab/meta-CLS.

## Appendix

**Figure A1.**
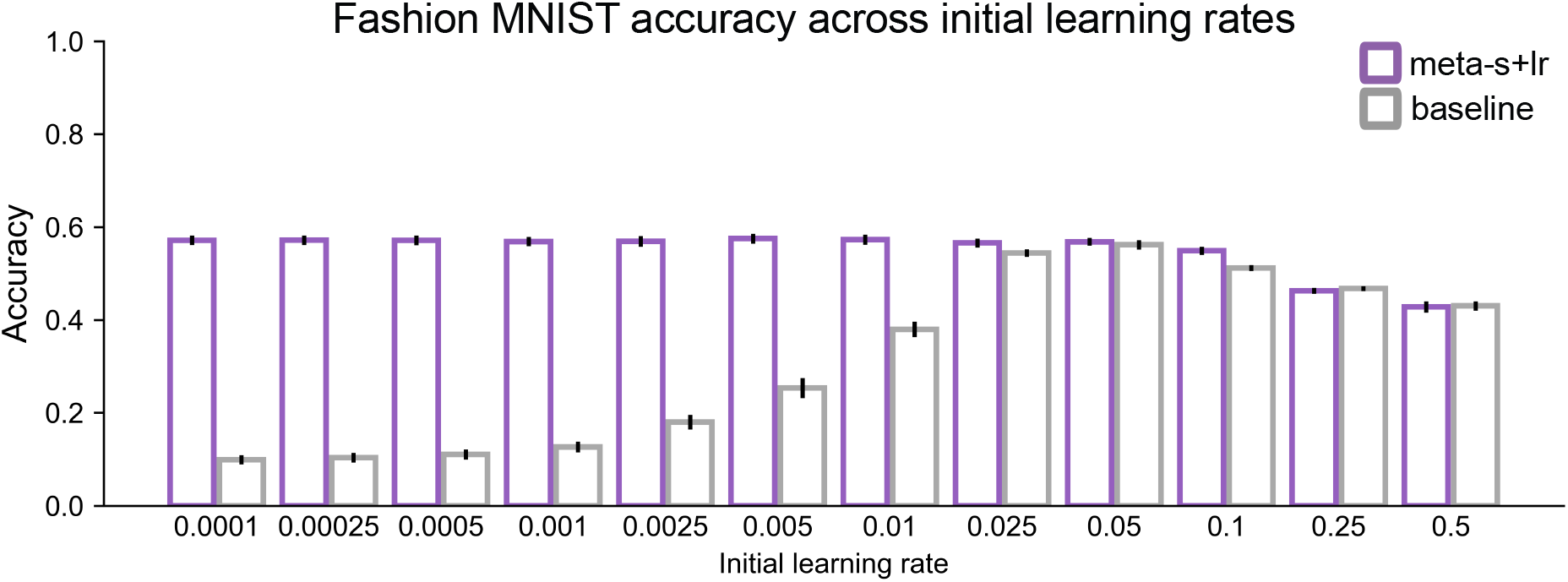
Test accuracy of meta-learned (green) vs baseline (orange) Fashion MNIST models across different initial learning rates.

**Figure A2.**
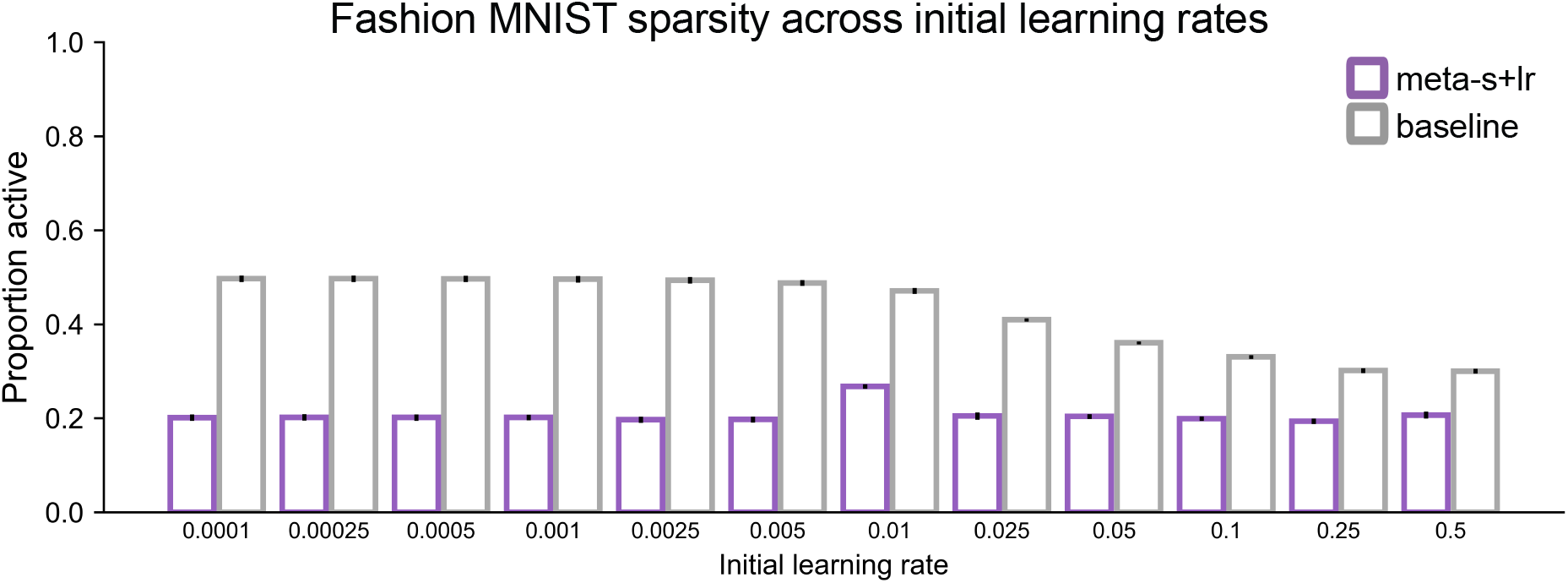
Proportion of hidden units active at test for meta-learned (green) and baseline (orange) Fashion MNIST models across different initial learning rates.

**Figure A3.**
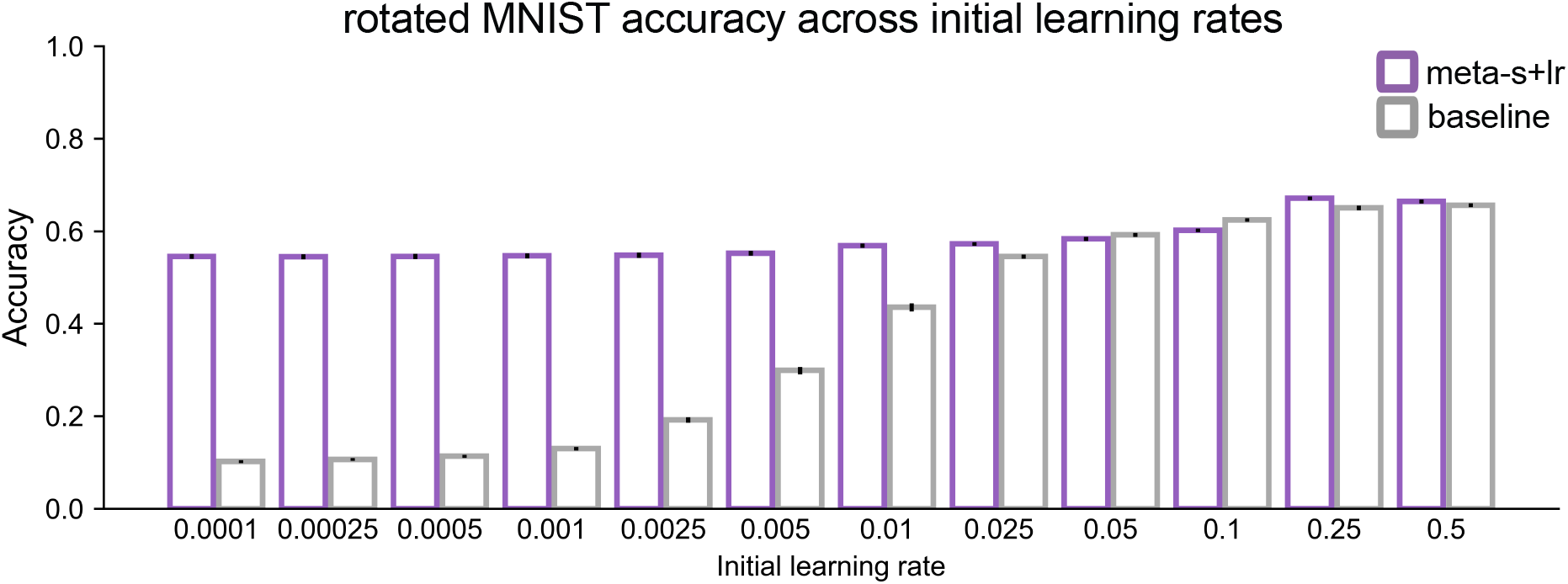
Test accuracy of meta-learned (green) vs baseline (orange) rotated MNIST models across initial learning rates.

**Figure A4.**
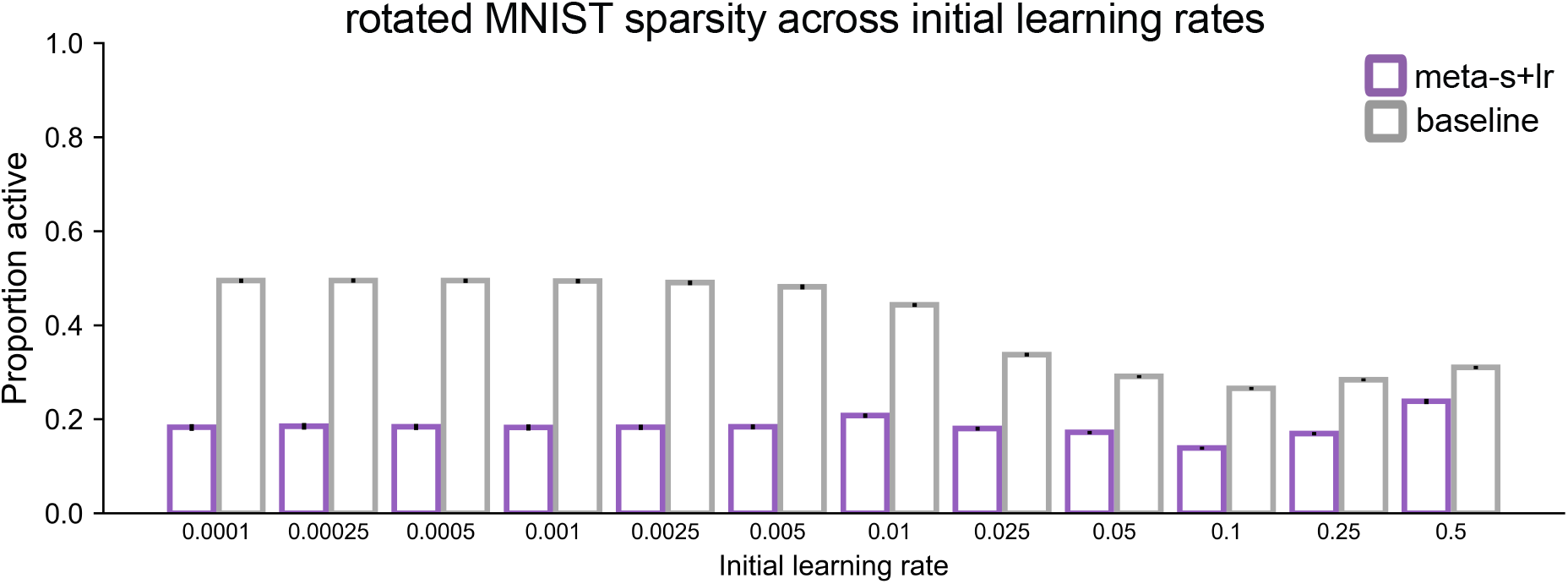
Proportion of hidden units active at test for meta-learned (green) and baseline (orange) rotated MNIST models across different initial learning rates.

**Figure A5.**
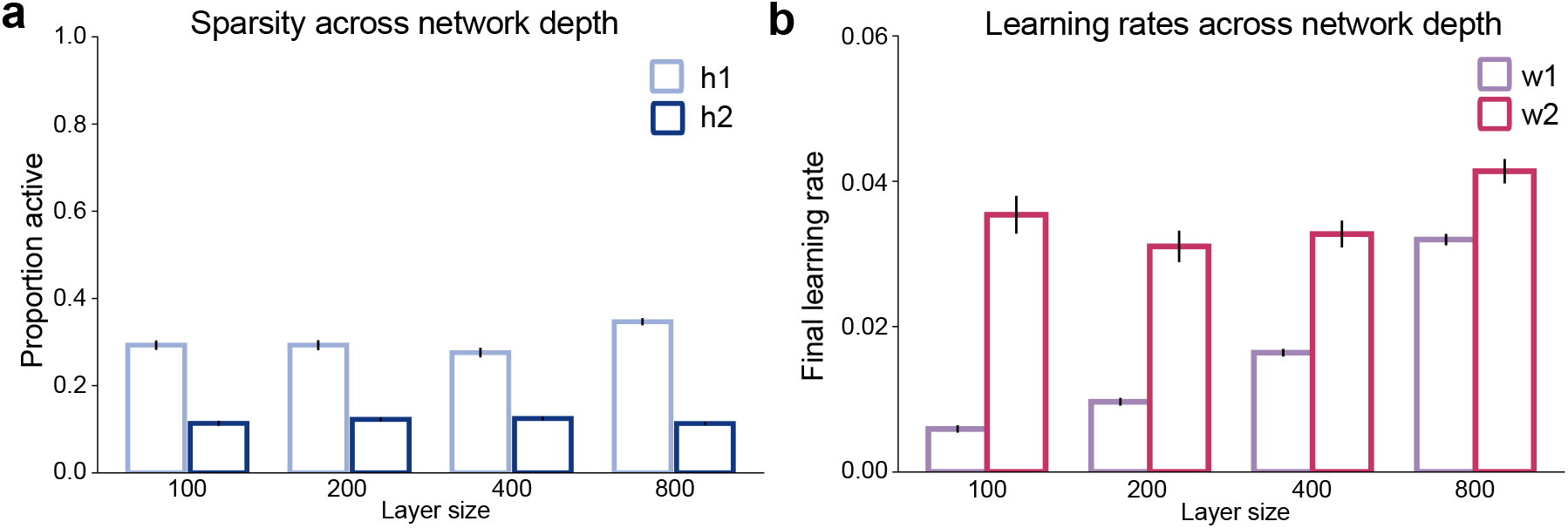
**a**. The mean proportion of units active per layer at test in shallow networks trained on rotated MNIST with varying hidden-layer sizes. **b**. Final learning rates for weights in shallow networks trained on rotated MNIST with hidden-layer sizes of 100, 200, 400, and 800 units; all other parameters were held constant.

**Figure A6.**
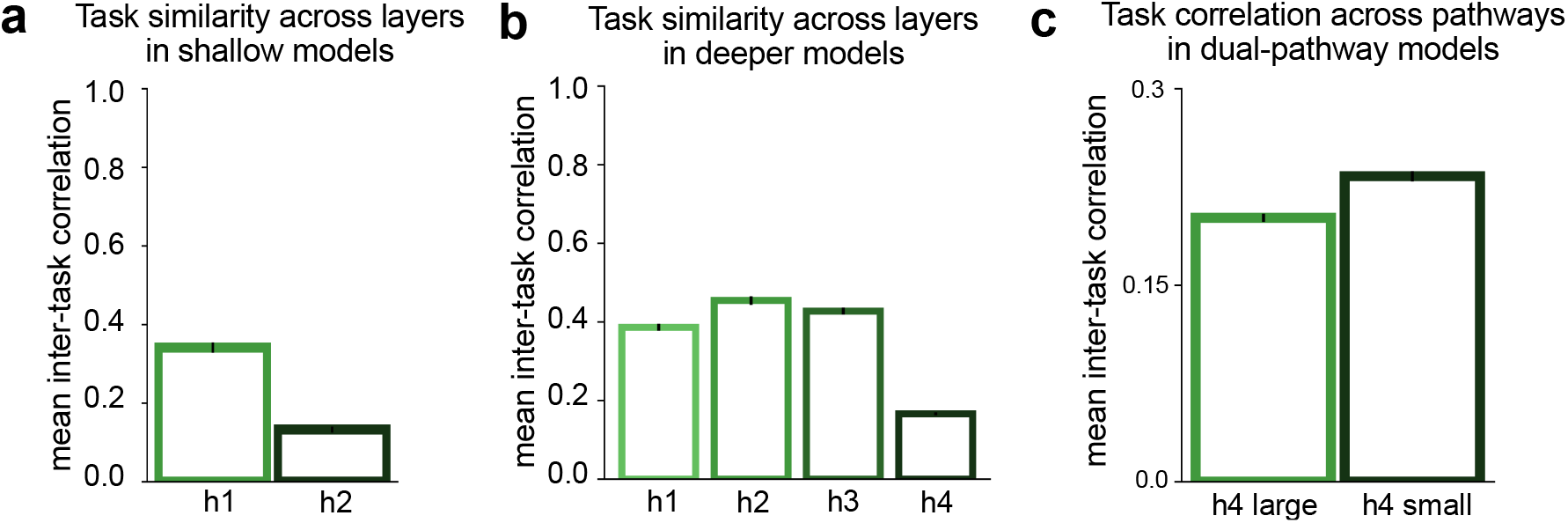
Mean inter-task representational correlation across layers (lower = stronger pattern separation).

## References

1. Felleman, D. J. and Essen, D. C. V. (1991). Distributed hierarchical processing in the primate cerebral cortex. Cerebral cortex, 1(1):1–47.

2. McClelland, J. L., McNaughton, B. L., and O’Reilly, R. C. (1995). Why there are complementary learning systems in the hippocampus and neocortex: insights from the successes and failures of connectionist models of learning and memory. Psychological review, 102(3):419.

3. Ison, M. J., Quian Quiroga, R., and Fried, I. (2015). Rapid encoding of new memories by individual neurons in the human brain. Neuron, 87(1):220–230.

4. Monaco, J. D., Rao, G., Roth, E. D., and Knierim, J. J. (2014). Attentive scanning behavior drives one-trial potentiation of hippocampal place fields. Nature Neuroscience, 17(5):725–731.

5. O’Reilly, R. C. and Rudy, J. W. (2001). Conjunctive representations in learning and memory: principles of cortical and hippocampal function. Psychological review, 108(2):311.

6. Rolls, E. T. and Treves, A. (2011). The neuronal encoding of information in the brain. Progress in Neurobiology, 95(3):448–490.

7. Yang, T. and Maunsell, J. H. R. (2004). The effect of perceptual learning on neuronal responses in monkey visual area v4. Journal of Neuroscience, 24(7):1617–1626.

8. Kobatake, E., Wang, G., and Tanaka, K. (1998). Effects of shape-discrimination training on the selectivity of inferotemporal cells in adult monkeys. Journal of neurophysiology, 80(1):324–330.

9. Ghose, G. M., Yang, T., and Maunsell, J. H. R. (2002). Physiological correlates of perceptual learning in monkey V1 and V2. Journal of neurophysiology, 87(4):1867–1888.

10. Miyashita, Y. (1988). Neuronal correlate of visual associative long-term memory in the primate temporal cortex. Nature, 335(6193):817–820.

11. Schoups, A., Vogels, R., Qian, N., and Orban, G. (2001). Practising orientation identification improves orientation coding in V1 neurons. Nature, 412(6846):549–553.

12. Quian Quiroga, R., Kraskov, A., Koch, C., and Fried, I. (2009). Explicit encoding of multimodal percepts by single neurons in the human brain. Current Biology, 19(15):1308–1313.

13. Reh, R. K., Dias, B. G., Nelson, 3rd, C. A., Kaufer, D., Werker, J. F., Kolb, B., Levine, J. D., and Hensch, T. K. (2020). Critical period regulation across multiple timescales. Proceedings of the National Academy of Sciences, 117(38):23242–23251.

14. Condé, F., Lund, J. S., and Lewis, D. A. (1996). The hierarchical development of monkey visual cortical regions as revealed by the maturation of parvalbumin-immunoreactive neurons. Brain Res. Dev. Brain Res., 96(1-2):261–276.

15. Quian Quiroga, R. (2012). Concept cells: the building blocks of declarative memory functions. Nature Reviews Neuroscience, 13(8):587–597.

16. Barnes, C. A., McNaughton, B. L., Mizumori, S. J. Y., Leonard, B. W., and Lin, L. H. (1990). Comparison of spatial and temporal characteristics of neuronal activity in sequential stages of hippocampal processing. 83:287–300.

17. Yassa, M. A. and Stark, C. E. L. (2011). Pattern separation in the hippocampus. Trends in Neuroscience, 34(10):515–525.

18. Mormann, F., Kornblith, S., Cerf, M., Ison, M. J., Kraskov, A., Tran, M., Knieling, S., Quian Quiroga, R., Koch, C., and Fried, I. (2017). Scene-selective coding by single neurons in the human parahippocampal cortex. Proceedings of the National Academy of Sciences, 114(5):1153–1158.

19. Leutgeb, S., Leutgeb, J. K., Treves, A., Moser, M.-B., and Moser, E. I. (2004). Distinct ensemble codes in hippocampal areas CA3 and CA1. Science, 305(5688):1295–1298.

20. Schapiro, A. C., Turk-Browne, N. B., Botvinick, M. M., and Norman, K. A. (2017). Complementary learning systems within the hippocampus: a neural network modelling approach to reconciling episodic memory with statistical learning. Philosophical Transactions of the Royal Society B: Biological Sciences, 372(1711):20160049.

21. Sun, W., Advani, M., Spruston, N., Saxe, A., and Fitzgerald, J. E. (2023). Organizing memories for generalization in complementary learning systems. Nature neuroscience, 26(8):1438–1448.

22. Whittington, J. C., Muller, T. H., Mark, S., Chen, G., Barry, C., and Burgess, N. (2020). The tolmaneichenbaum machine: unifying space and relational memory through generalization in the hippocampal formation. Cell, 183(5):1249–1263.

23. Spens, E. and Burgess, N. (2024). A generative model of memory construction and consolidation. Nature Human Behaviour, pages 1–18.

24. Singh, D., Norman, K. A., and Schapiro, A. C. (2022). A model of autonomous interactions between hippocampus and neocortex driving sleep-dependent memory consolidation. Proceedings of the National Academy of Sciences, 119(44):e2123432119.

25. Geerts, J. P., Chersi, F., Stachenfeld, K. L., and Burgess, N. (2020). A general model of hippocampal and dorsal striatal learning and decision making. Proceedings of the National Academy of Sciences, 117(49):31427–31437.

26. Remme, M. W. H., Bergmann, U., Alevi, D., Schreiber, S., Sprekeler, H., and Kempter, R. (2021). Hebbian plasticity in parallel synaptic pathways: A circuit mechanism for systems memory consolidation. PLoS Comput. Biol., 17(12):e1009681.

27. Marr, D., Willshaw, D., and McNaughton, B. (1991). Simple memory: a theory for archicortex. In From the Retina to the Neocortex, pages 59–128. Birkhäuser Boston.

28. Sherry, D. F. and Schacter, D. L. (1987). The evolution of multiple memory systems. Psychological Review, 94(4):439–454.

29. Kumaran, D., Hassabis, D., and McClelland, J. L. (2016). What learning systems do intelligent agents need? complementary learning systems theory updated. Trends in Cognitive Sciences, 20(7):512–534.

30. Norman, K. A. and O’Reilly, R. C. (2003). Modeling hippocampal and neocortical contributions to recognition memory: a complementary-learning-systems approach. Psychological Review, 110(4):611–646.

31. Fusi, S., Drew, P. J., and Abbott, L. F. (2005). Cascade models of synaptically stored memories. Neuron, 45(4):599–611.

32. Ba, J., Hinton, G. E., Mnih, V., Leibo, J. Z., and Ionescu, C. (2016). Using fast weights to attend to the recent past. In Advances in Neural Information Processing Systems, volume 29.

33. Goldstein, A., Zada, Z., Buchnik, E., Schain, M., Price, A., Aubrey, B., and Hasson, U. (2022). Shared computational principles for language processing in humans and deep language models. Nature Neuroscience, 25(3):369–380.

34. Khaligh-Razavi, S.-M. and Kriegeskorte, N. (2014). Deep supervised, but not unsupervised, models may explain it cortical representation. PLoS Computational Biology, 10(11):e1003915.

35. Schrimpf, M., Blank, I. A., Tuckute, G., Kauf, C., Hosseini, E. A., Kanwisher, N., Tenenbaum, J. B., and Fedorenko, E. (2021). The neural architecture of language: Integrative modeling converges on predictive processing. Proceedings of the National Academy of Sciences, 118(45).

36. Yamins, D. L. K., Hong, H., Cadieu, C. F., Solomon, E. A., Seibert, D., and DiCarlo, J. J. (2014). Performance-optimized hierarchical models predict neural responses in higher visual cortex. Proceedings of the national academy of sciences, 111(23):8619–8624.

37. Gupta, G., Yadav, K., and Paull, L. (2020). Look-ahead meta learning for continual learning. Advances in Neural Information Processing Systems, 33:11588–11598.

38. Duchi, J., Hazan, E., and Singer, Y. (2011). Adaptive subgradient methods for online learning and stochastic optimization. Journal of Machine Learning Research, 12(7).

39. Kingma, D. P. (2014). Adam: A Method for Stochastic Optimization.

40. Bergstra, J. and Bengio, Y. (2012). Random search for hyper-parameter optimization. Journal of Machine Learning Research, 13(1):281–305.

41. Li, Z., Zhou, F., Chen, F., and Li, H. (2017). Meta-SGD: Learning to Learn Quickly for Few-Shot Learning.

42. Bricken, T., Davies, X., Singh, D., Krotov, D., and Kreiman, G. (2023). Sparse distributed memory is a continual learner. ICLR.

43. Ahmad, S. and Scheinkman, L. (2019). How can we be so dense? the benefits of using highly sparse representations. arXiv, abs/1903.11257.

44. Fedus, W., Zoph, B., and Shazeer, N. (2022). Switch transformers: Scaling to trillion parameter models with simple and efficient sparsity. Journal of Machine Learning Research, 23(120):1–39.

45. Makhzani, A. and Frey, B. (2014). k-sparse autoencoders. CoRR, abs/1312.5663.

46. Rumelhart, D. and Zipser, D. (1985). Feature discovery by competitive learning. Cognitive Science, 9(1):75–112.

47. Matthew Riemer, Ignacio Cases, R. A. M. L. I. R. Y. T. and Tesauro, G. (2019). Learning to learn without forgetting by maximizing transfer and minimizing interference. International Conference on Learning Representations.

48. Kirkpatrick, J., Pascanu, R., Rabinowitz, N., Veness, J., Desjardins, G., Rusu, A. A., Milan, K., Quan, J., Ramalho, T., Grabska-Barwinska, A., Hassabis, D., Clopath, C., Kumaran, D., and Hadsell, R. (2017). Overcoming catastrophic forgetting in neural networks. Proceedings of the National Academy of Sciences, 114(13):3521–3526.

49. Lopez-Paz, D. and Ranzato, M. (2017). Gradient episodic memory for continual learning. Advances in neural information processing systems, page 6467–6476.

50. Li, Z., Zhou, F., Chen, F.,, and Li, H. (2017). Meta-sgd: Learning to learn quickly for few-shot learning. arXiv.

51. Hoefler, T., Alistarh, D., Ben-Nun, T., Dryden, N., and Peste, A. (2021). Sparsity in deep learning: Pruning and growth for efficient inference and training in neural networks. The Journal of Machine Learning Research, 22(1):10882–11005.

52. Wixted, J. T., Squire, L. R., Jang, Y., Papesh, M. H., Goldinger, S. D., Kuhn, J. R., Smith, K. A., Treiman, D. M., and Steinmetz, P. N. (2014). Sparse and distributed coding of episodic memory in neurons of the human hippocampus. Proceedings of the National Academy of Sciences, 111(26):9621–9626.

53. Olshausen, B. A. and Field, D. J. (1996). Emergence of simple-cell receptive field properties by learning a sparse code for natural images. Nature, 381(6583):607–609.

54. Javed, K. and White, M. (2019). Meta-learning representations for continual learning. Advances in Neural Information Processing Systems, 32.

55. Baker, S., Vieweg, P., Gao, F., Gilboa, A., Wolbers, T., Black, S. E., and Rosenbaum, R. S. (2016). The human dentate gyrus plays a necessary role in discriminating new memories. Current Biology, 26(19):2629– 2634.

56. Lee, I., Rao, G., and Knierim, J. J. (2004). A double dissociation between hippocampal subfields: differential time course of CA3 and CA1 place cells for processing changed environments. Neuron, 42(5):803–815.

57. Leutgeb, S., Leutgeb, J. K., Moser, E. I., and Moser, M. B. (2006). Fast rate coding in hippocampal ca3 cell ensembles. Hippocampus, 16(9):765–774.

58. van Dijk, M. T. and Fenton, A. A. (2018). On how the dentate gyrus contributes to memory discrimination. Neuron, 98(4):832–845.

59. Hainmueller, T. and Bartos, M. (2020). Dentate gyrus circuits for encoding, retrieval and discrimination of episodic memories. Nature Reviews Neuroscience, 21(3):153–168.

60. Wang, H. S., Rosenbaum, R. S., Baker, S., Lauzon, C., Batterink, L. J., and köhler, S. (2023). Dentate gyrus integrity is necessary for behavioral pattern separation but not statistical learning. Journal of Cognitive Neuroscience, 35(5):900–917.

61. Ketz, N., Morkonda, S. G., and O’Reilly, R. C. (2013). Theta coordinated error-driven learning in the hippocampus. PLoS Computational Biology, 9(6):e1003067.

62. Zhou, Z., Singh, D., Tandoc, M. C., and Schapiro, A. C. (2023). Building integrated representations through interleaved learning. Journal of Experimental Psychology: General, 152(9):2666–2684.

63. Sun, Y., Nguyen, A. Q., Nguyen, J. P., Le, L., Saur, D., Choi, J., Callaway, E. M., and Xu, X. (2014). Cell-type-specific circuit connectivity of hippocampal ca1 revealed through cre-dependent rabies tracing. Cell Reports, 7(1):269–280.

64. Witter, M. P., Wouterlood, F. G., Naber, P. A., and Van Haeften, T. (2006). Anatomical organization of the parahippocampal-hippocampal network. Annals of the New York Academy of Sciences, 911(1):1–24.

65. Sučević, J. and Schapiro, A. C. (2023). A neural network model of hippocampal contributions to category learning. eLife, 12:e77185.

66. Kang, L. and Toyoizumi, T. (2024). Distinguishing examples while building concepts in hippocampal and artificial networks. Nature Communications, 15(1):647.

67. Crist, R. E., Li, W., and Gilbert, C. D. (2001). Learning to see: experience and attention in primary visual cortex. Nature Neuroscience, 4(5):519–525.

68. Naya, Y., Yoshida, M., and Miyashita, Y. (2001). Backward spreading of memory-retrieval signal in the primate temporal cortex. Science, 291(5504):661–664.

69. Cadena, S. A., Willeke, K. F., Restivo, K., Denfield, G., Sinz, F. H., Bethge, M., Tolias, A. S., and Ecker, A. S. (2024). Diverse task-driven modeling of macaque V4 reveals functional specialization towards semantic tasks. PLoS Computational Biology, 20(5):e1012056.

70. Rust, N. C. and DiCarlo, J. J. (2012). Balanced increases in selectivity and tolerance produce constant sparseness along the ventral visual stream. Journal of Neuroscience, 32(30):10170–10182.

71. Kumaran, D. and McClelland, J. L. (2012). Generalization through the recurrent interaction of episodic memories: a model of the hippocampal system. Psychological review, 119(3):573.

72. Ning, W., Bladon, J. H., and Hasselmo, M. E. (2022). Complementary representations of time in the prefrontal cortex and hippocampus. Hippocampus, 32(8):577–596.

73. Barlow, H. B. (1972). Single units and sensation: a neuron doctrine for perceptual psychology? Perception, 1(4):371–394.

74. Kent, B. A., Hvoslef-Eide, M., Saksida, L. M., and Bussey, T. J. (2016). The representational–hierarchical view of pattern separation: Not just hippocampus, not just space, not just memory? Neurobiology of learning and memory, 129:99–106.

75. Solomon, S., Kay, K., and Schapiro, A. (2024). Recent statistics shift object representations in parahippocampal cortex. bioRxiv, 2024-02.

76. O’Reilly, R. C. and Norman, K. A. (2002). Hippocampal and neocortical contributions to memory: Advances in the complementary learning systems framework. Trends in cognitive sciences, 6(12):505–510.

77. Ahissar, M. and Hochstein, S. (2004). The reverse hierarchy theory of visual perceptual learning. Trends in cognitive sciences, 8(10):457–464.

78. Hinton, G. E. (1984). Distributed representations. Technical Report CMU-CS-84-157, Carnegie-Mellon University, Computer Science Department.

79. Chini, M., Pfeffer, T., and Hanganu-Opatz, I. (2022). An increase of inhibition drives the developmental decorrelation of neural activity. eLife, 11:e78811.

80. Xin, W., Kaneko, M., Roth, R. H., Zhang, A., Nocera, S., Ding, J. B., Stryker, M. P., and Chan, J. R. (2024). Oligodendrocytes and myelin limit neuronal plasticity in visual cortex. Nature, 633(8031):856–863.

81. McGee, A. W., Yang, Y., Fischer, Q. S., Daw, N. W., and Strittmatter, S. M. (2005). Experience-driven plasticity of visual cortex limited by myelin and Nogo receptor. Science, 309(5744):2222–2226.

82. Pinto, J. G., Hornby, K. R., Jones, D. G., and Murphy, K. M. (2010). Developmental changes in gabaergic mechanisms in human visual cortex across the lifespan. Frontiers in Cellular Neuroscience, 4:1421.

83. Porges, E. C., Jensen, G., Foster, B., Edden, R. A., and Puts, N. A. (2021). The trajectory of cortical gaba across the lifespan, an individual participant data meta-analysis of edited mrs studies. eLife, 10:e62575.

84. Perica, M. I., Calabro, F. J., Larsen, B., Foran, W., Yushmanov, V. E., Hetherington, H., and Luna, B. (2022). Development of frontal gaba and glutamate supports excitation/inhibition balance from adolescence into adulthood. Progress in Neurobiology, 219:102370.

85. Baum, G. L., Flournoy, J. C., Glasser, M. F., Harms, M. P., Mair, P., Sanders, A. F. P., Barch, D. M., et al. (2022). Graded variation in t1w/t2w ratio during adolescence: measurement, caveats, and implications for development of cortical myelin. Journal of Neuroscience, 42(29):5681–5694.

86. Grydeland, H., Vertes, P. E., Vasa, F., Romero-Garcia, R., Whitaker, K., Alexander-Bloch, A. F., Bjornerud, A., et al. (2019). Waves of maturation and senescence in micro-structural mri markers of human cortical myelination over the lifespan. Cerebral Cortex, 29(3):1369–1381.

87. Zhu, Y., Sousa, A. M. M., Gao, T., Skarica, M., Li, M., Santpere, G., and Esteller-Cucala, P. (2018). Spatiotemporal transcriptomic divergence across human and macaque brain development. Science, 362(6420):eaat8077.

88. Miller, D. J., Duka, T., Stimpson, C. D., Schapiro, S. J., Baze, W. B., McArthur, M. J., and Fobbs, J. (2012). Prolonged myelination in human neocortical evolution. Proceedings of the National Academy of Sciences, 109(41):16480–16485.

89. de Faria Jr., O., Pivonkova, H., Varga, B., Timmler, S., Evans, K. A., and Káradóttir, R. T. (2021). Periods of synchronized myelin changes shape brain function and plasticity. Nature Neuroscience, 24(11):1508– 1521.

90. Nussenbaum, K. and Hartley, C. A. (2024). Understanding the development of reward learning through the lens of meta-learning. Nature Review Psychology.

91. Schwartz, R., Dodge, J., Smith, N. A., and Etzioni, O. (2020). Green ai. Communications of the ACM, 63(12):54–63.

92. Olshausen, B. and Field, D. (2004). Sparse coding of sensory inputs. Current Opinion in Neurobiology, 14(4):481–487.

93. Klinzing, J. G., Niethard, N., and Born, J. (2019). Mechanisms of systems memory consolidation during sleep. Nature Neuroscience, 22(10):1598–1610.

